# De novo genome assembly of an interspecific hybrid grapevine ‘Maeve’

**DOI:** 10.1101/2025.03.24.644857

**Authors:** Manabu Fujie, Mayumi Kawamitsu, Hidetoshi Saze

## Abstract

The grapevine is one of the most ancient and economically important horticultural crops in the world. The grapevine species *Vitis vinifera* is cultivated globally; however, due to its susceptibility to pathogens and environmental stresses, wild *Vitis* species and their hybrids are often used as rootstocks in vineyards. Here, we report the genome analysis of a *Vitis* strain named Maeve, which was identified in a vineyard in Japan, though its genetic origin remains unclear. We performed haplotype-resolved de novo assembly of the Maeve genome using PacBio HiFi sequencing and Hi-C assembly. Our genome analysis revealed that Maeve has originated from an interspecific hybridization between an unknown *V. vinifera* cultivar and the *V. riparia* Gloire cultivar, likely arising through breeding or natural pollination. This novel cultivar has a potential to expand wine grape production across a wider range of environmental conditions where conventional cultivars are unsuitable for viticulture.

## Introduction

The grape (*Vitis* spp.) is one of the most widely cultivated horticultural crops globally, primarily used for wine production and as table fruit (This et al. 2006; Zhou et al. 2017). Due to relatively weak reproductive barriers among Vitis species and their close wild relatives, interspecific hybrids are frequently bred and propagated (Myles et al. 2011; Dong et al. 2023). These hybrids facilitate the introgression of beneficial traits from wild grape species into cultivated varieties, making them valuable genetic resources for agriculture. With advancements in high-throughput and long-read sequencing technologies, an increasing number of Vitis genomes—encompassing both cultivated and wild species—have been sequenced, including haplotype-resolved whole-genome assemblies (Liang et al. 2019; Cochetel et al. 2023; Liu et al. 2024; Guo et al. 2025).

The European grape (*Vitis vinifera* L.) is predominantly grown for wine and juice production (Shi et al. 2023; Velt et al. 2023). However, due to its susceptibility to various biotic and abiotic stresses, it is commonly grafted onto rootstocks in vineyards to enhance resilience. These rootstocks, mainly derived from North American wild grape species or their interspecific hybrids, provide resistance to pests such as phylloxera and lime, as well as tolerance to environmental challenges, including drought and cold stress (Cochetel et al. 2023; Cantu et al. 2024). Commonly used rootstock varieties include the riverbank grape *V. riparia* (e.g., Gloire de Montpellier) and its hybrids with *V. berlandieri* (e.g., 5BB, 5C, SO4) or *V. rupestris* (e.g., 101- 14, 3309)(Minio et al. 2022). Additionally, hybrids between *V. rupestris* and *V. berlandieri* (e.g., 110R, 1103P) are widely utilized. Beyond wine grapes, interspecific hybridization has also contributed to the development of table grape varieties, such as Shine Muscat, which has a breeding lineage involving *V. labruscana* and *V. vinifera* (Shirasawa et al. 2022). These examples underscore the significance of wild grape species as genetic reservoirs for enhancing agronomic and economic traits in grapevine breeding.

A red grapevine cultivar named “Maeve” was discovered in a vineyard in Kanagawa, Japan (Tanaka and Ishimura 2020). This cultivar exhibits remarkable adaptability and productivity across Japan’s diverse climates, ranging from the subarctic conditions of Hokkaido to the subtropical environment of Okinawa. Recent studies have indicated that Maeve possesses superior freezing tolerance compared to standard *V. vinifera* cultivars (Kita et al. 2024). Although prior genetic analysis using microsatellite markers suggested that Maeve may have *V. riparia* ancestry (Tanaka and Ishimura 2020), its exact contribution remains uncertain. In this study, we conducted a haplotype-resolved *de novo* assembly of the Maeve genome using PacBio HiFi sequencing and Hi-C assembly. Our genomic analysis revealed that Maeve originated from an interspecific hybridization between an unidentified *V. vinifera* cultivar and the *V. riparia* Gloire cultivar. This unique lineage suggests that Maeve could contribute to expanding wine grape production in regions where conventional grapevine cultivation is currently unfeasible.

## Materials and Methods

### High molecular DNA extraction, library construction and sequencing

Plant samples were collected from the Komesu Grape Farm in Nago, Japan. For PacBio HiFi sequencing, high-molecular-weight (HMW) genomic DNA was extracted by isolating nuclei from young Maeve leaves that had been disrupted in liquid nitrogen. The extraction was performed using the Nanobind® HMW DNA kit (PacBio), following the standard protocol for plant nuclei. A HiFi sequencing library was then constructed using the SMRTbell® prep kit 3.0 (PacBio). During this process, genomic DNA was sheared into fragments with a peak size of 25 kb using the Megaruptor 3 system (Diagenode). Fragments smaller than 17 kb were removed using BluePippin (High Pass Plus™ Gel Cassettes, BPLUS).

Sequencing was carried out on the PacBio Sequel II platform using an SMRT Cell 8M tray (PacBio; 101-389-001), along with the Binding Kit 3.2 and cleanup beads (PacBio; 102-333-300) and Sequel II sequencing kit 2.0(101-820-200). Additionally, Revio sequencing was performed using a Revio SMRT Cell tray (PacBio; 102-202-200) and the Revio™ Polymerase Kit (PacBio; 102-739-100) and Revio sequencing plate(102-587-400). Data yields are summarized in Table S1.

### Hi-C library preparation and sequencing

Prior to nuclei isolation for Hi-C library preparation, mature leaves were incubated in a dark container at 20°C for 66 hours while being supplied with water through the petiole. Crosslinking was performed following the protocol provided in the Arima Hi-C kit (Arima Genomics). Nuclei were extracted from frozen leaf tissue using the CelLytic™ PN Isolation/Extraction Kit (Sigma-Aldrich) according to the manufacturer’s protocol for obtaining highly purified nuclei.

The Hi-C library was then prepared using the Dovetail Omni-C Kit (Dovetail Genomics). Sequencing was conducted on the NovaSeq X Plus platform using the NovaSeq™ X Series 1.5B Reagent Kit (300 Cycles). Data yields are reported in Table S1.

### Genome assembly

To estimate genome homozygosity and heterozygosity, k-mer analysis was performed using Jellyfish (Marçais and Kingsford 2011) and GenomeScope (Vurture et al. 2017). Hi-C Illumina reads were preprocessed using fastp (Chen et al. 2018) with default settings. Initial genome assembly was carried out with PacBio Revio HiFi reads and Hi-C Illumina reads using Hifiasm v0.19.9 (Cheng et al. 2021) with the following parameters: -l 0, --telo-m CCCTAAA, --hom-cov 205, based on k-mer analysis.

Contigs were aligned to the plastid and mitochondrial genomes of Vitis vinifera PN40024.v4 (Velt et al. 2023) using Minimap2 v2.28-r1209 (Li 2018). Nine contigs that showed over 30% identity to organelle genomes were excluded from the final assembly. Hi-C data was mapped onto the assembled contigs using Juicer v1.6.2 (Durand et al. 2016), and a Hi-C contact map was generated with 3D-DNA v180922 (Dudchenko et al. 2017). Chromosome structures were reviewed in Juicebox v1.11.08 (https://github.com/aidenlab/Juicebox), and manual scaffold assembly and corrections were applied.

To close assembly gaps, PacBio Sequel II HiFi reads were processed using TGS-GapCloser (Xu et al. 2020) with the parameters --ne --tgstype pb --minmap_arg -x asm20. Finally, scaffolds smaller than 2 Mb were removed, resulting in a total of 38 scaffolds (Table 1). Alignment against the PN40024 reference genome was performed and visualized using D-GENIES (Cabanettes and Klopp 2018).

**Table 1.**
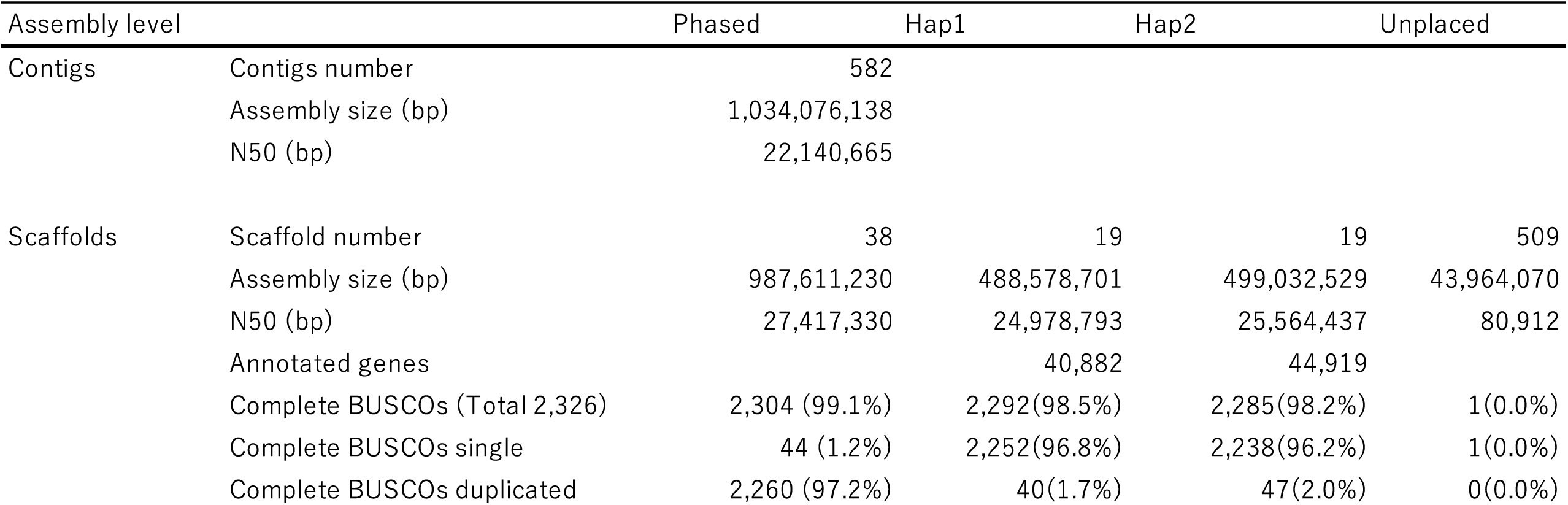
Assembly statistics and BUSCO scores of the whole, haplotype 1, and haplotype 2 of Maeve genome assembly in this study.

### Repeat and gene annotations

The assembled genome was anchored to the Vitis vinifera PN40024 reference genome (Shi et al. 2023), along with genomes of wild Vitis species and interspecific hybrids (Minio et al. 2022; Cochetel et al. 2023), using Minimap2 v2.28-r1209 (Li 2018) with default settings. Haploid chromosomes were classified based on sequence similarity to *V. vinifera* and *V. riparia* genomes (see details in the main text).

Repeat and gene annotations were performed separately for each haploid genome. Repeats were identified using Extensive de novo TE Annotator (EDTA) v2.2.2 (Ou et al. 2019) with a Vitis repeat library downloaded from (https://www.grapegenomics.com/pages/repeats.php). RepeatMasker v4.1.5 (https://repeatmasker.org) was applied to soft-mask the genome sequences with the parameters -q -gff -html -a -u -xsmall.

Gene prediction was conducted using BRAKER3 (Gabriel et al. 2024 Feb) with default settings. The annotation was supported by a comprehensive peptide library, combining sequences from: The plant peptide library (Viridiplantae.fa) from OrthoDB 11 (https://bioinf.uni-greifswald.de/bioinf/partitioned_odb11/), The Vitis vinifera peptide library (Vitis_vinifera.PN40024.v4.pep.all.fa) from EnsemblPlants (https://plants.ensembl.org/Vitis_vinifera/), The Vitis riparia peptide library (VITVri588271_v1.0.protein.fasta) from Grapegenomics.com (https://www.grapegenomics.com). Functional gene annotation was conducted using Hayai-Annotation Plants v2.0 (https://pgdbjsnp.kazusa.or.jp/app/hayai2), based on amino acid sequences generated from the BRAKER3 pipeline (braker.aa).

Transfer RNAs (tRNAs) were identified using tRNAscan-SE v2.0.12 (Chan et al. 2021). Ribosomal RNA (rRNA), microRNA (miRNA), and small nucleolar RNA (snoRNA) were detected using INFERNAL v1.1.5 (Nawrocki and Eddy 2013) with the Rfam.cm database.

Plastid and mitochondrial genomes were assembled using PacBio Revio HiFi reads with Oatk (Zhou et al. 2024) and the Oatk embryophyta database. The mitochondrial genome assembly was further refined and scaffolded using the Vitis vinifera PN40024.v4 mitochondrial genome (https://plants.ensembl.org/Vitis_vinifera/) as a reference, employing Ragtag v2.1.0 (Alonge et al. 2022).

### Genome similarity analysis between Vitis haplotypes

To assess sequence similarity between Maeve and other Vitis strains, Maeve chromosomes were aligned against publicly available Vitis and interspecific hybrid genomes (Minio et al. 2022; Cochetel et al. 2023) as well as the Vitis pan-genome dataset (Guo et al. 2025) using MUMmer4 v4.0.0rc1 (Marçais et al. 2018). The percentage of aligned bases was used as a value for genome similarity. Heatmaps for similarity matrices were generated using the ComplexHeatmap package in R (Gu et al. 2016).

For structural variation analysis, genome alignment paf files between Maeve haplotypes and other Vitis species were generated using Minimap2 v2.28-r1209 (Li 2018) with the parameters -x asm20 -c -eqx -secondary=no. Structural variations were visualized using SVbyEye v0.99 in R (Porubsky et al. 2024) with the following settings: perc.identity.breaks = c(85, 90, 95), min.query.aligned.bp = 8000000. Genome synteny between Maeve and Vitis species was analyzed using ntSynt (Coombe et al. 2024) with the parameter -d 10 and visualized using ntSynt_viz (Coombe et al. 2025).

## Results

### Haplotype-Resolved Genome Assembly of Maeve

To investigate the genetic background and genome structure of the grapevine cultivar Maeve (Figure 1), we performed a haploid-resolved *de novo* genome assembly using PacBio Sequel II and Revio HiFi reads, achieving >70× and >99× diploid genome coverage, respectively (Table S1). The initial assembly yielded 582 contigs with a total genome size of 1,034,076,138 bp (Table 1). Further refinement using Hi-C Illumina sequencing reads resulted in 38 major scaffolds, which exhibited high-frequency contact patterns (Figure 2A), consistent with the diploid chromosome number of Vitis species (Velt et al. 2023).

**Figure 1.**
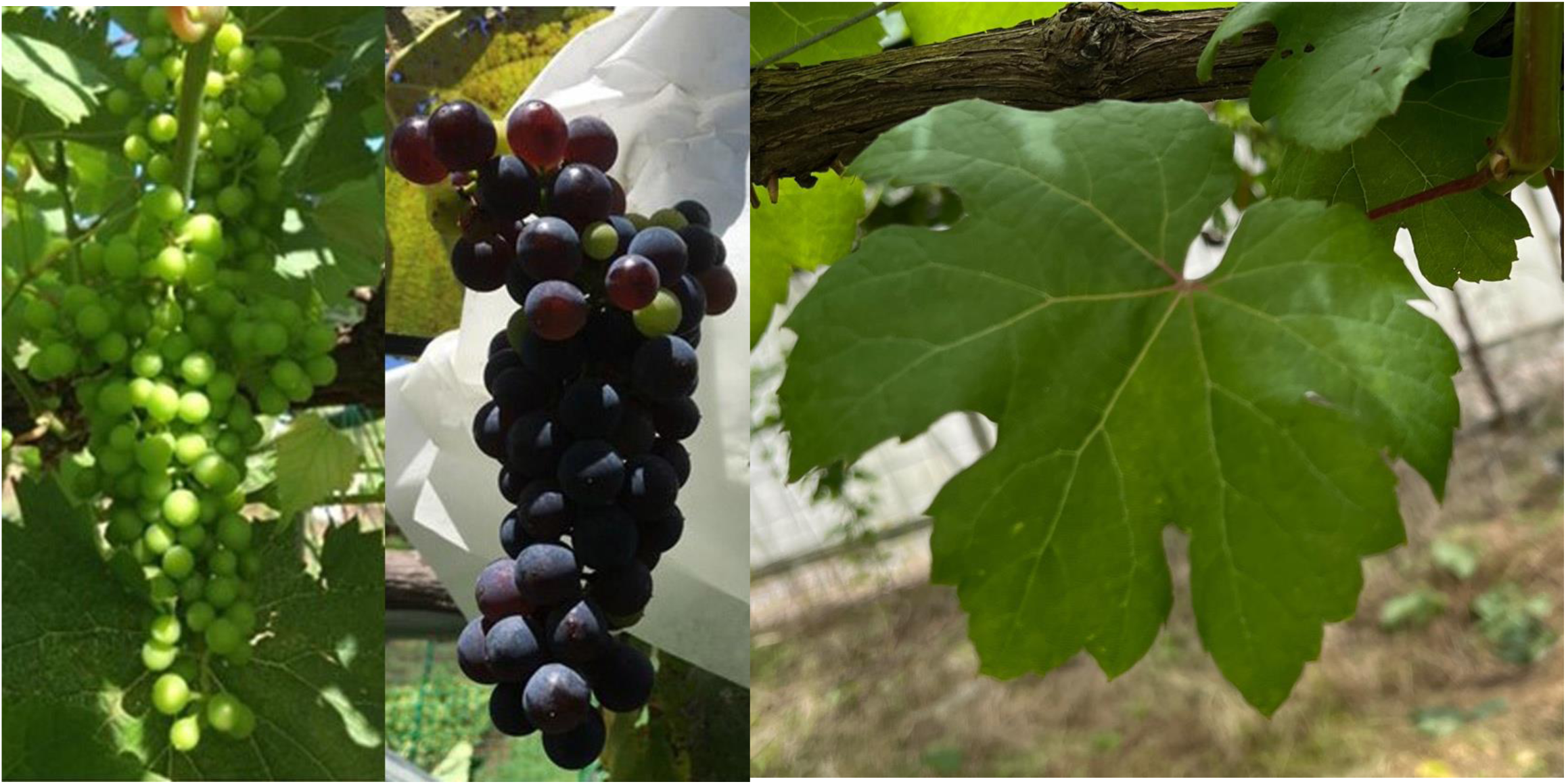
The Vitis variety ‘Maeve’. Young (left) and mature (middle) berries and leaf (right) of the Maeve cultivar.

**Figure 2.**
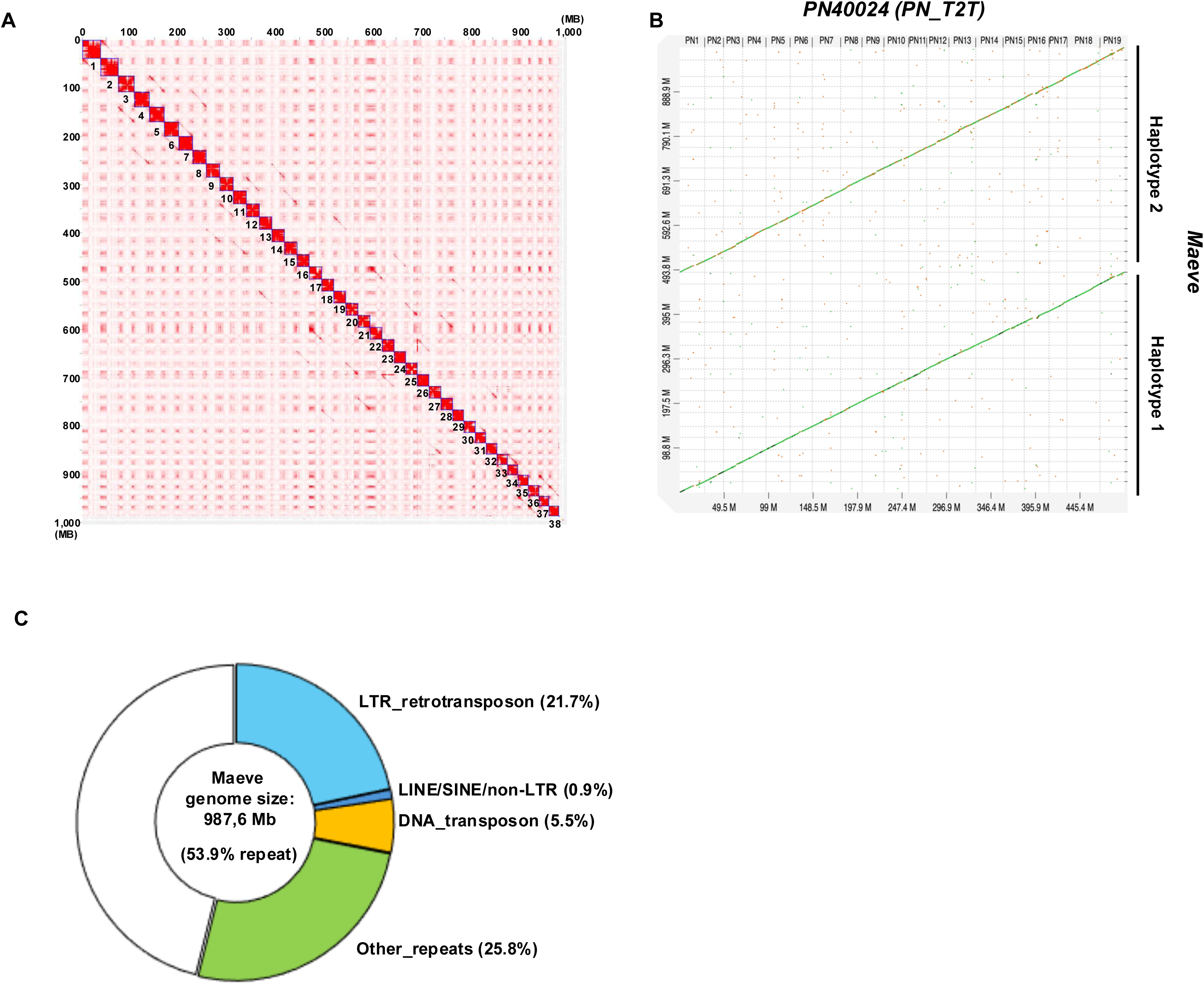
The haplotype-resolved genome assembly of Maeve. (A) Hi-C contact map of Maeve assembly. The numbers in the matrix indicate independent scaffolds. (B) Alignment of the phased Maeve genome assembly against the reference PN40024 genome (). (C) Repeat content of the Maeve genome assembly.

Alignment of the assembled scaffolds against the *V. vinifera* PN40024 reference genome (PN_T2T)(Shi et al. 2023) revealed collinearity between Maeve’s two haplotype chromosomes and the *V. vinifera* genome, though with variable sequence similarity (Figure 2B). Based on their similarity to the *V. vinifera* PN40024 genome, the haplotypes were classified as Haplotype 1 and Haplotype 2 (Table S2; see Materials and Methods). Haplotype 1 spanned 488,578,701 bp, while haplotype 2 spanned 499,032,529 bp. BUSCO analysis indicated high completeness scores for the whole genome (99.1%), as well as for each haplotype (98.5% for haplotype 1 and 98.2% for haplotype 2) (Table 1).

Annotation of repetitive sequences revealed that 53.9% of the genome consisted of repeats, with long terminal repeat (LTR) retrotransposons being the most abundant class (21.7%) (Figure 2C, Table S3). Gene prediction identified 40,882 genes in haplotype 1 and 44,919 genes in haplotype 2 (Table 1, Supplementary Figure S1, Tables S4–S6).

### High Heterozygosity in the Maeve Genome

To assess the structural variation and sequence divergence between the two haplotypes, we performed a self-alignment of the homologous assemblies. The results revealed relatively low sequence similarity between haplotype 1 and haplotype 2 (Figure 3A), indicating a genetic divergence between the two haplotypes and minimal recombination due to inbreeding. K-mer analysis (*k*=21) of the Maeve genome further supported its high heterozygosity, estimating a heterozygosity rate of 2.99% (estimated by GenomeScope)(Vurture et al. 2017), in contrast to the reference *V. vinifera* Pinot Noir genome (PN40024)(Shi et al. 2023), which exhibited a much lower heterozygosity of 0.115%. These findings suggest that Maeve originated from a hybridization event involving genetically divergent parental lineages.

**Figure 3.**
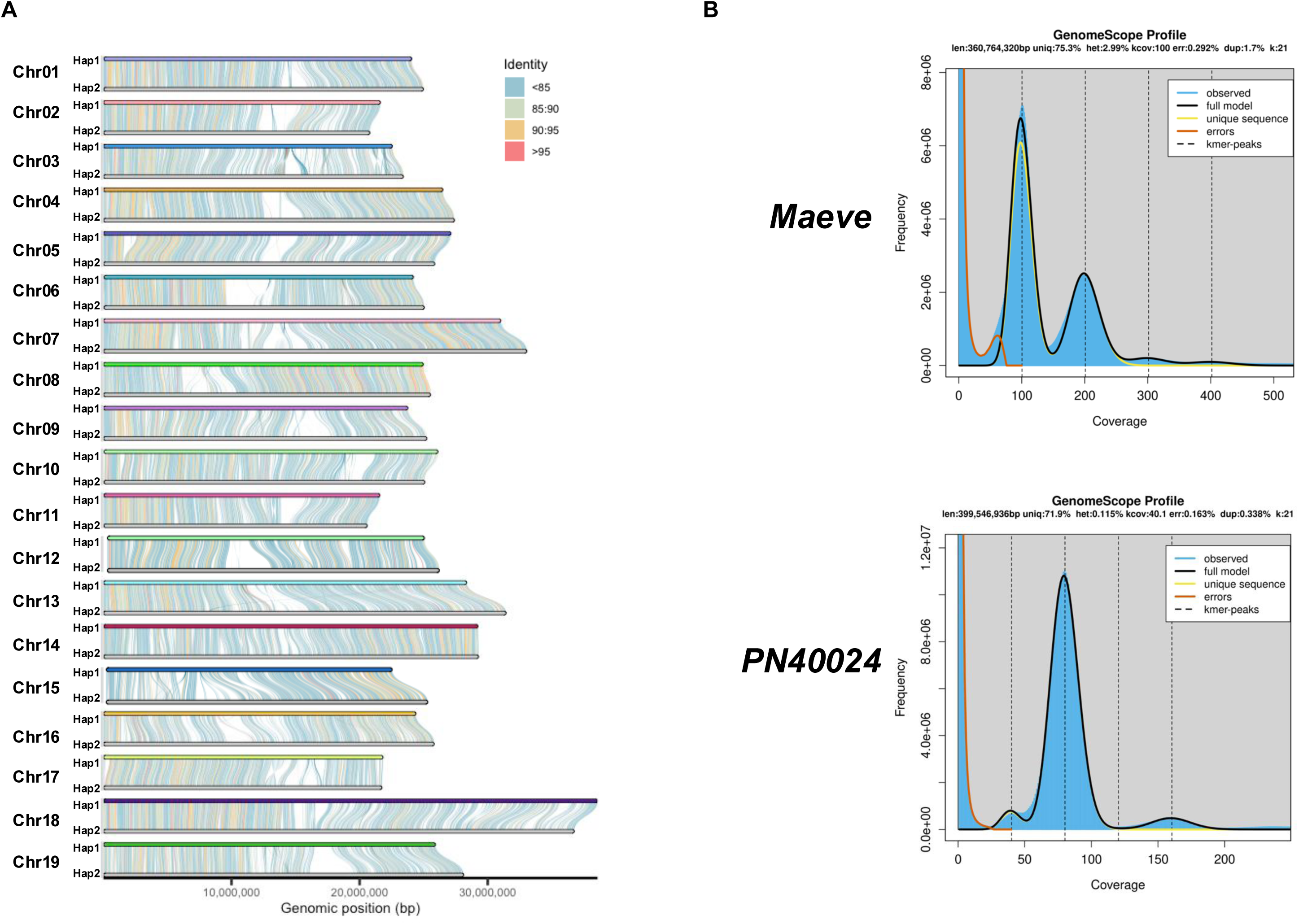
Heterozygosity of the Maeve genome. (A) Structural synteny and similarity between Hap1 and Hap2 of the Maeve genome assembly. (B) K-mer coverage and frequency (k= 21) estimated in the Maeve genome sequence (upper) and the reference PN40024 genome sequence (lower).

### Maeve Genome Indicates an Interspecific Hybrid Origin

The pronounced divergence between Maeve’s haplotypes suggests that its genome may contain chromosomal introgressions from other Vitis species. Previous studies have proposed an interspecific origin for Maeve, specifically involving contributions from the wild grape *Vitis riparia* (Tanaka and Ishimura 2020). To investigate Maeve’s genomic ancestry, we aligned its haplotype 1 and haplotype 2 sequences to publicly available genome assemblies of *V. vinifera* (Shi et al. 2023) and several wild Vitis species, including *V. berlandieri*, *V. rupestris*, and *V. riparia* (Cochetel et al. 2023). Additionally, we compared Maeve’s genome with sequences of interspecific hybrid cultivars commonly used as rootstocks, such as 101-14 Mgt (*V. riparia* × *V. rupestris*) and Kober 5BB (*V. riparia* × *V. berlandieri*) (Minio et al. 2022). The results revealed distinct genomic similarities between Maeve’s haplotypes and different Vitis species (Figure 4A, B). Haplotype 1 showed the highest similarity to the *V. vinifera* reference genome (Figure 4A, B), whereas haplotype 2 exhibited greater similarity to Kober 5BB, an interspecific hybrid of *V. riparia* × *V. berlandieri* (Figure 4A, C). Notably, Maeve haplotype 2 was particularly similar to the *V. riparia*-derived haplotype of Kober 5BB, strongly indicating that Maeve is derived from a cross between *V. vinifera* and *V. riparia* or a closely related species.

**Figure 4.**
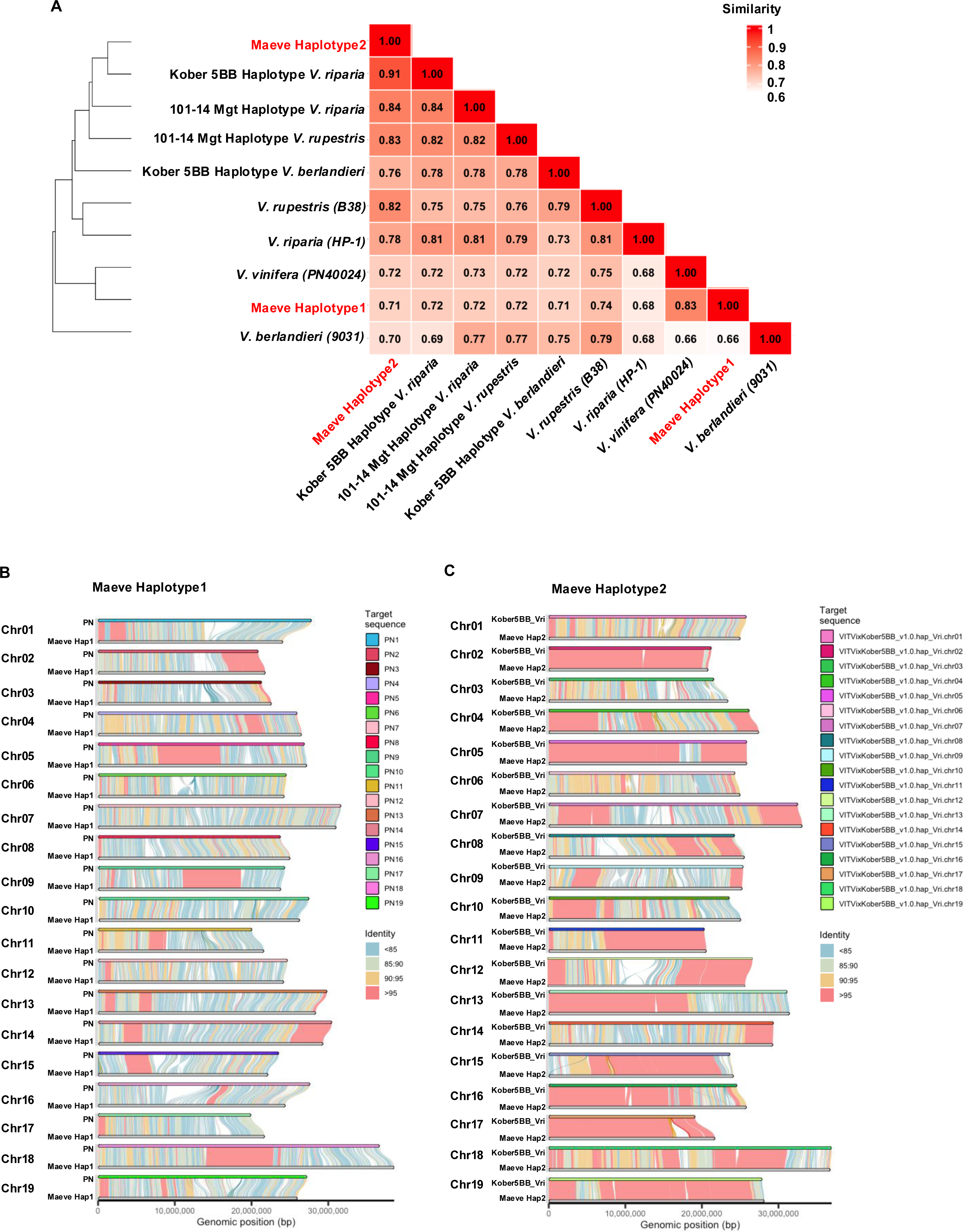
Maeve genome originates from interspecific hybridization of Vitis species. (A) A triangle heatmap showing seminaries between Maeve haploid genome assemblies, haploid genomes of wild Vitis species, and interspecies hybrid. Color gradients indicate similarity between two assembly (0.6 to 1). (B) Comparison of the Maeve haplotype 1 (Hap1) and the reference PN40024 (PN). Genome regions showing similarities were connected by blocks and identities of the genomes were indicated as color maps. (C) Comparison of the Maeve haplotype 2 (Hap2) and the haplotype *V. riparia* of Kober 5BB as (B).

### Maeve Originated from an Interspecific Cross Between *V. vinifera* and *V. riparia*

To further resolve Maeve’s ancestry, we compared its genome against a recently published dataset comprising 142 haplotypes from 71 Vitis genomes (Guo et al. 2025), including both wild and cultivated varieties (Table S7). The analysis demonstrated that Maeve haplotype 1 shares ∼92.77% sequence similarity with multiple *V. vinifera* cultivars, including Riesling and Chardonnay (Figure 5A, C, S4). However, it did not exhibit perfect identity with any specific cultivar, suggesting an unidentified *V. vinifera* parent. In contrast, Maeve haplotype 2 showed the highest similarity to both haplotypes of *V. riparia* cultivar Gloire (V048) (∼99.85%; Figure 5B, D). Further examination of chromosomal segments revealed that Maeve haplotype 2 inherited complementary sequences from both haplotypes of *V. riparia* Gloire, indicating that recombination occurred between these parental haplotypes (Figure S5). This confirms that one of Maeve’s parental strains is *V. riparia* Gloire, further supporting its interspecific hybrid origin.

**Figure 5.**
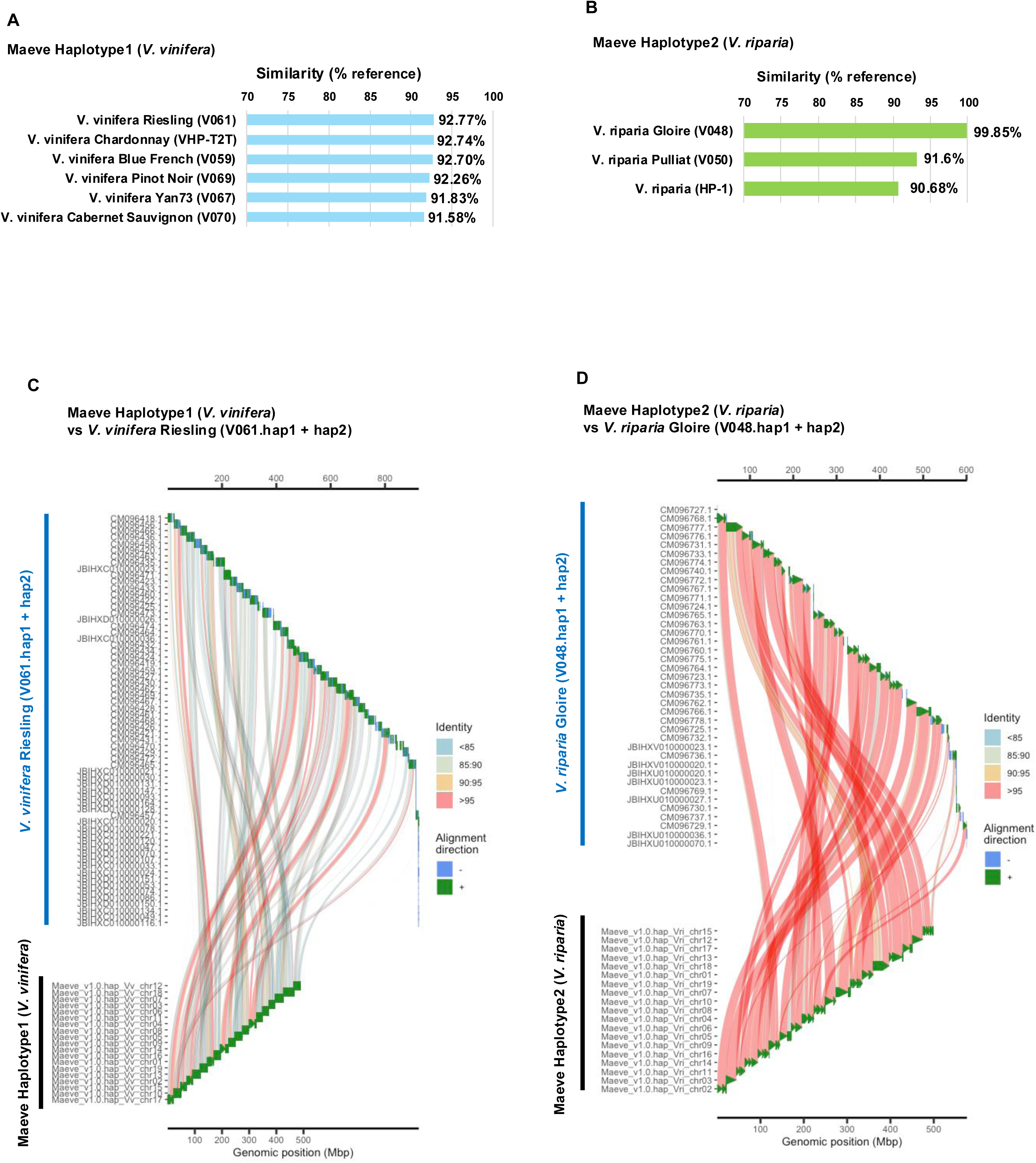
Comparison of Maeve haplotypes to Vitis species. (A) Similarity (aligned sequence % of the Maeve haplotype 1; *V. vinifera* origin) to the indicated *V. vinifera* cultivars (haplotype 1 and haplotype 2). (B) Similarity (aligned sequence % of the Maeve haplotype 2; *V. riparia* origin) to the indicated *V. riparia* cultivars (haplotype 1 and haplotype 2). (C) An alignment of Maeve haplotype 1 and *V. vinifera* Riesling chromosomes (haplotype 1 and haplotype 2). Sequence identities were shown in the color codes as indicated in the legend. (D) An alignment of Maeve haplotype 2 and *V. riparia* Gloire chromosomes (haplotype 1 and haplotype 2). Sequence identities were shown in the color codes as indicated in the legend.

## Discussion

In this study, we conducted a haplotype-resolved *de novo* assembly of the Maeve genome using PacBio HiFi sequencing and Hi-C technology. Our genome analysis revealed that Maeve originated from an interspecific hybridization event between an unidentified *V. vinifera* cultivar and *V. riparia* Gloire (Figure 5, S5). This hybridization may have occurred either through intentional breeding efforts or as a result of natural cross-pollination in the vineyard.

Hybridization between *V. vinifera* and *V. riparia* is not uncommon, as several cultivars have been developed from similar genetic backgrounds. For example, the widely used rootstock cultivar Castel 196-17 was derived from a cross between a *V. vinifera* cultivar and *V. riparia* Gloire de Montpellier (de Andrés et al. 2007). Likewise, wine grape cultivars such as Marquette and Baco Noir are known to have inherited genetic material from both *V. vinifera* and *V. riparia* through their breeding histories (Hemstad and Luby 2008; Balint and Reynolds 2013; Frioni et al. 2017). These cultivars have been selected for their enhanced resistance to environmental stresses and improved adaptability, characteristics that are also observed in Maeve. However, whether Maeve shares any direct genetic lineage with these cultivars remains unclear due to a lack of historical records tracking its introduction into Japanese vineyards. Future genomic comparisons could provide insights into the possible connections between Maeve and other interspecific hybrids.

One of the most striking traits of Maeve is its high adaptability to a wide range of climatic conditions, from the subarctic climate of Hokkaido to the subtropical environment of Okinawa (Tanaka 2022; Kita et al. 2024). This broad environmental tolerance suggests that Maeve has inherited key stress-resistance traits from *V. riparia*, which is known for its cold hardiness and resilience to various abiotic stresses (Cochetel et al. 2023; Cantu et al. 2024). In particular, previous studies have reported that Maeve exhibits significantly higher freezing tolerance than standard *V. vinifera* cultivars (Kita et al. 2024), a trait likely inherited from *V. riparia*. This genetic contribution could make Maeve a valuable resource for viticulture in regions prone to harsh winters, potentially expanding the geographical range of wine grape production. Beyond cold tolerance, *V. riparia* is also recognized for its resistance to several biotic stresses, including fungal pathogens and pests such as phylloxera. While the extent of Maeve’s resistance to these challenges remains to be fully characterized, its genetic background suggests that it may possess traits that could contribute to more sustainable viticulture with reduced reliance on chemical treatments. Future studies should evaluate Maeve’s resistance to common grapevine diseases and pests, which pose major threats to vineyard productivity.

Although our genomic analysis successfully identified *V. riparia* Gloire as one of Maeve’s parental strains (Figure 5, S5), the exact identity of the *V. vinifera* parent remains unresolved. Among the 142 haplotypes from 71 Vitis genomes examined in this study, no single *V. vinifera* cultivar showed perfect identity to Maeve haplotype 1. However, it exhibited the highest similarity to multiple *V. vinifera* cultivars, including Riesling and Chardonnay, suggesting that its *V. vinifera* ancestor may be closely related to these varieties. The lack of a precise match highlights the need for further expansion of genomic databases, which would improve our ability to trace the ancestry of interspecific hybrids like Maeve. Moreover, transcriptomic and functional studies could help elucidate how specific genetic variations contribute to Maeve’s unique agronomic traits.

Our findings revealed that Maeve as a distinct interspecific hybrid with a genetic composition derived from both *V. vinifera* and *V. riparia*. Future research should focus on further characterizing its resilience to environmental stressors, identifying its exact *V. vinifera* ancestry, and exploring its potential contributions to sustainable viticulture. Maeve represents a promising cultivar that could help expand grape production into regions where traditional *V. vinifera* cultivars struggle to thrive, ultimately broadening the scope of viticulture in the face of climate change.

### Author contributions

HS designed the research. MF, NK performed genome DNA extraction, sequence library preparation and sequencing analysis. HS analyzed the sequencing data and wrote the manuscript.

### Data availability

Raw PacBio Hifi sequence reads and Hi-C sequence reads have been submitted to NCBI Sequence Read Archive under Bioproject PRJNA1231776.

The genome assembly sequence and annotations of Maeve accessible via the following weblink: https://www.oist.jp/research/research-units/peu/genome-data.

## Acknowledgements

This work was supported by Shonan Corp. and OIST. We thank Komesu grape farm for sample collection of Maeve, and OIST SQC for PacBio Hifi sequencing and Hi-C Illumina sequencing. We thank Drs. Yutaka Jitsuyama, Hiroto Homma for valuable discussion and inputs to this study.

---

**Supplementary Figure 1.**
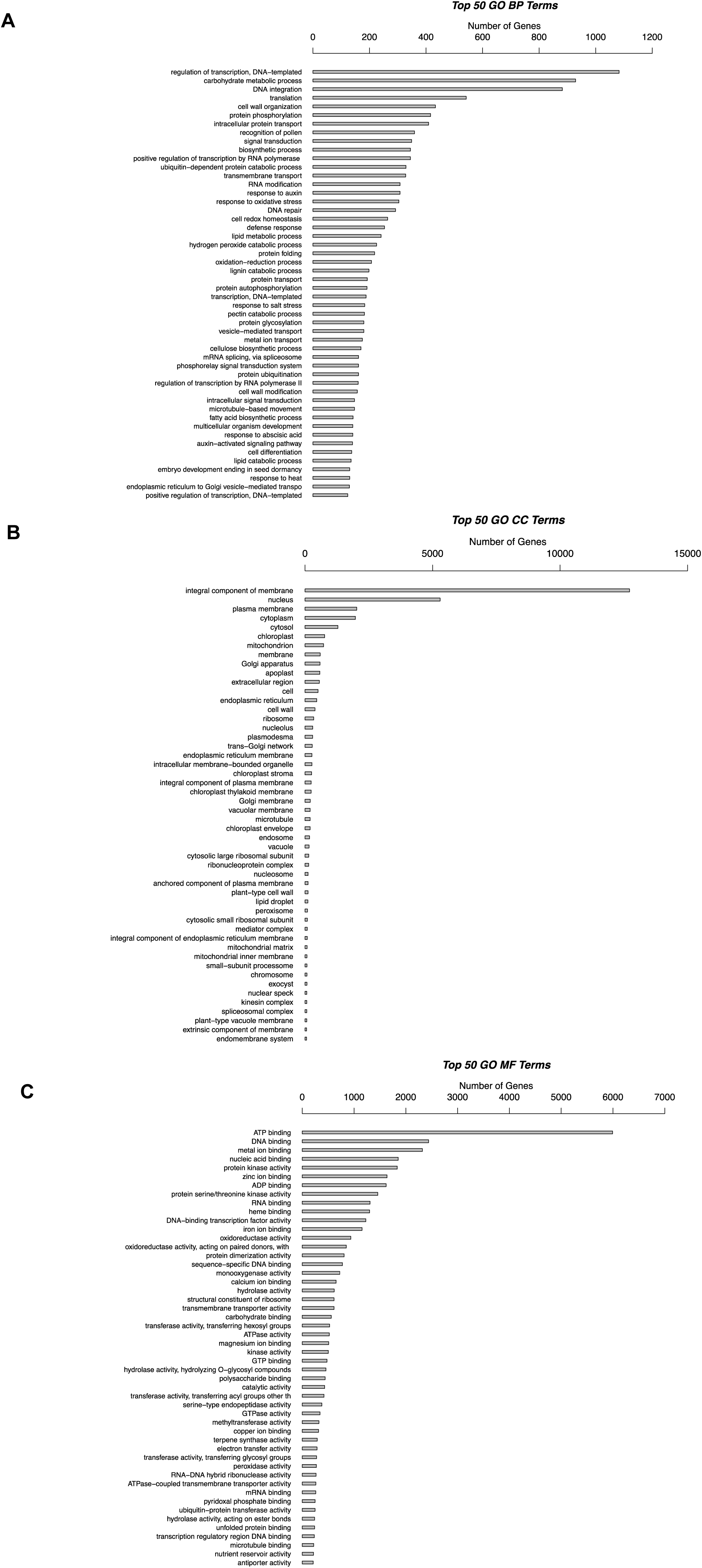
Gene Ontology (GO) of Maeve. (A) Top 50 GO terms for Biological Process (BP) and the number of annotated gene. (B) Top 50 GO terms for Cellular Component (CC) and the number of annotated genes. (C) Top 50 GO terms for Molecular Function (MF) and the number of annotated genes.

**Supplementary Figure 2.**
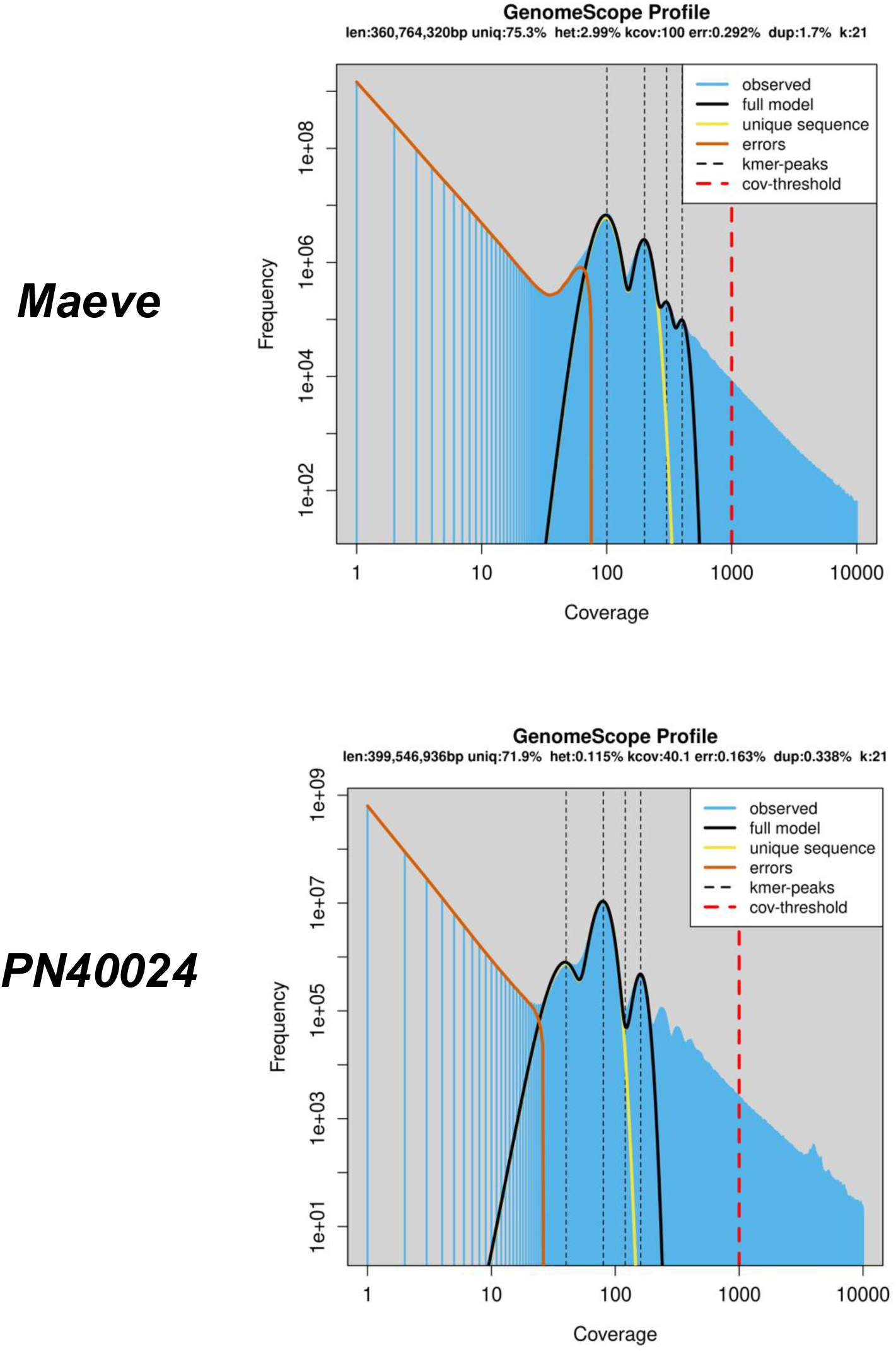
Heterozygosity of the Maeve genome. K-mer coverage and frequency (k= 21) estimated in the Maeve genome sequence (upper) and the reference PN40024 genome sequence (lower) as Fig. 3B, shown in log10 scale in x-axis.

**Supplementary Figure 3.**
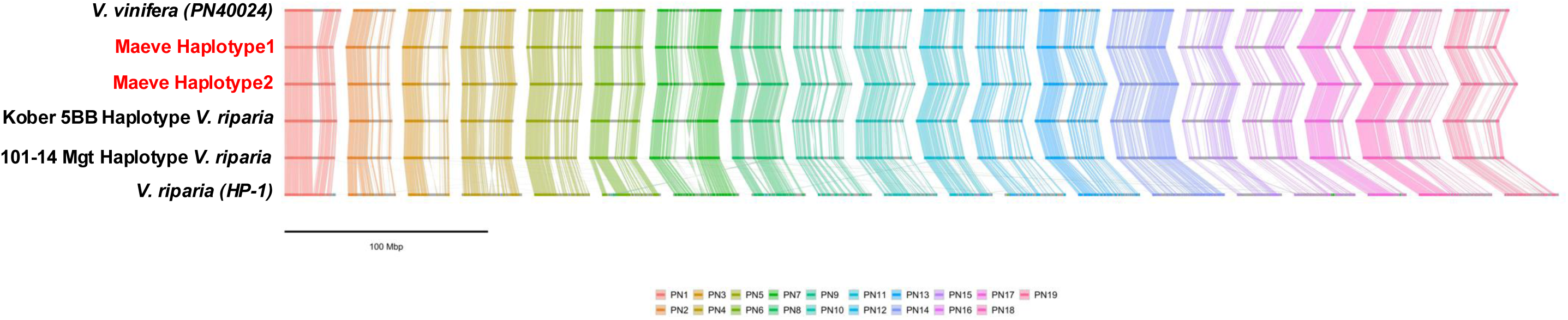
Synteny of Vitis spp. Genomes. Synteny blocks in chromosomes of *Vitis* species and interspecies hybrids including haplotype 1 and haplotype 2 of the Maeve genome.

**Supplementary Figure 4.**
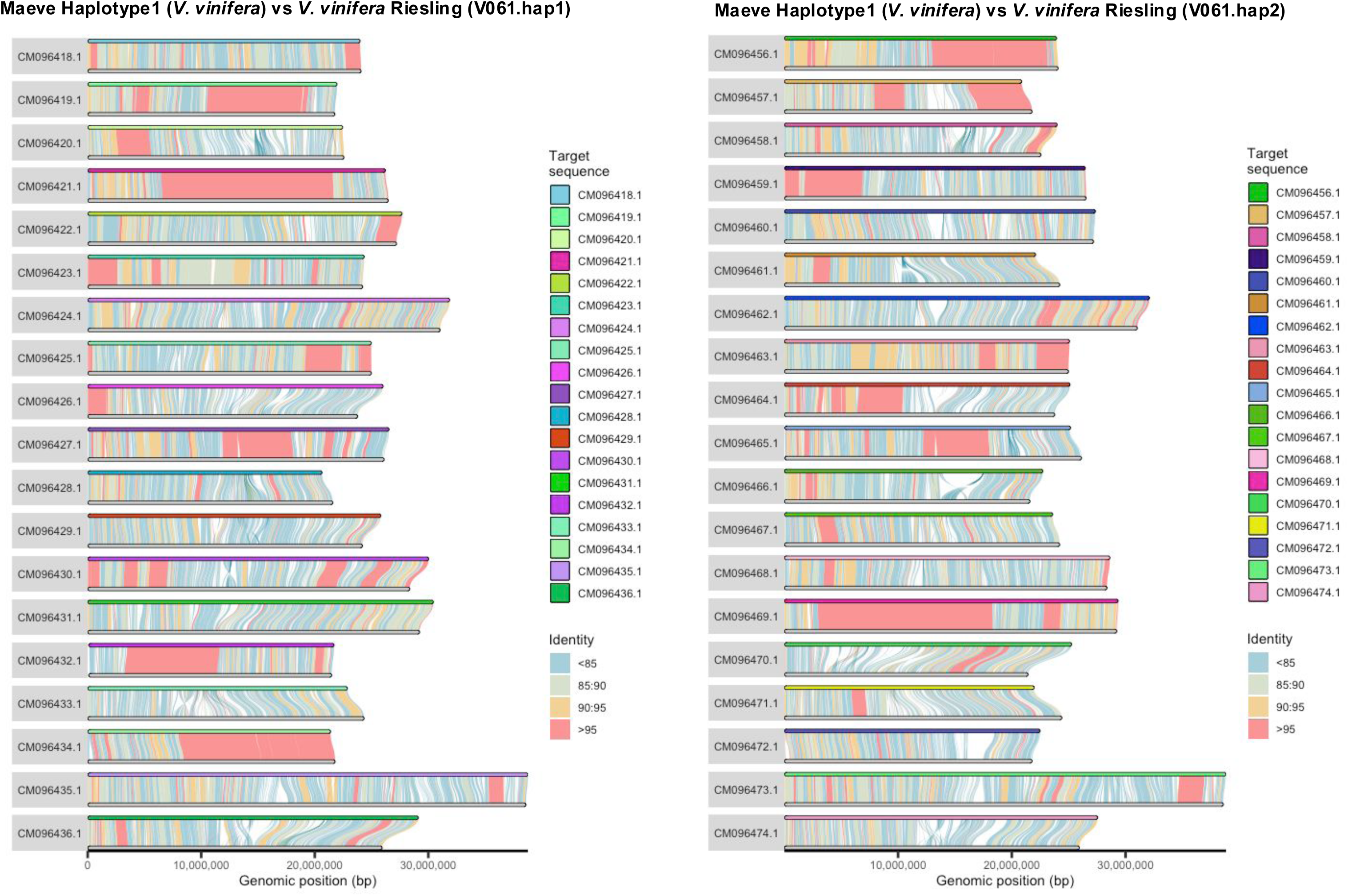
Synteny of Maeve haplotype 1 and Vitis vinifera genomes. Synteny blocks in chromosomes of *V. vinifera* Riesling and haplotype 1 of the Maeve genome.

**Supplementary Figure 5.**
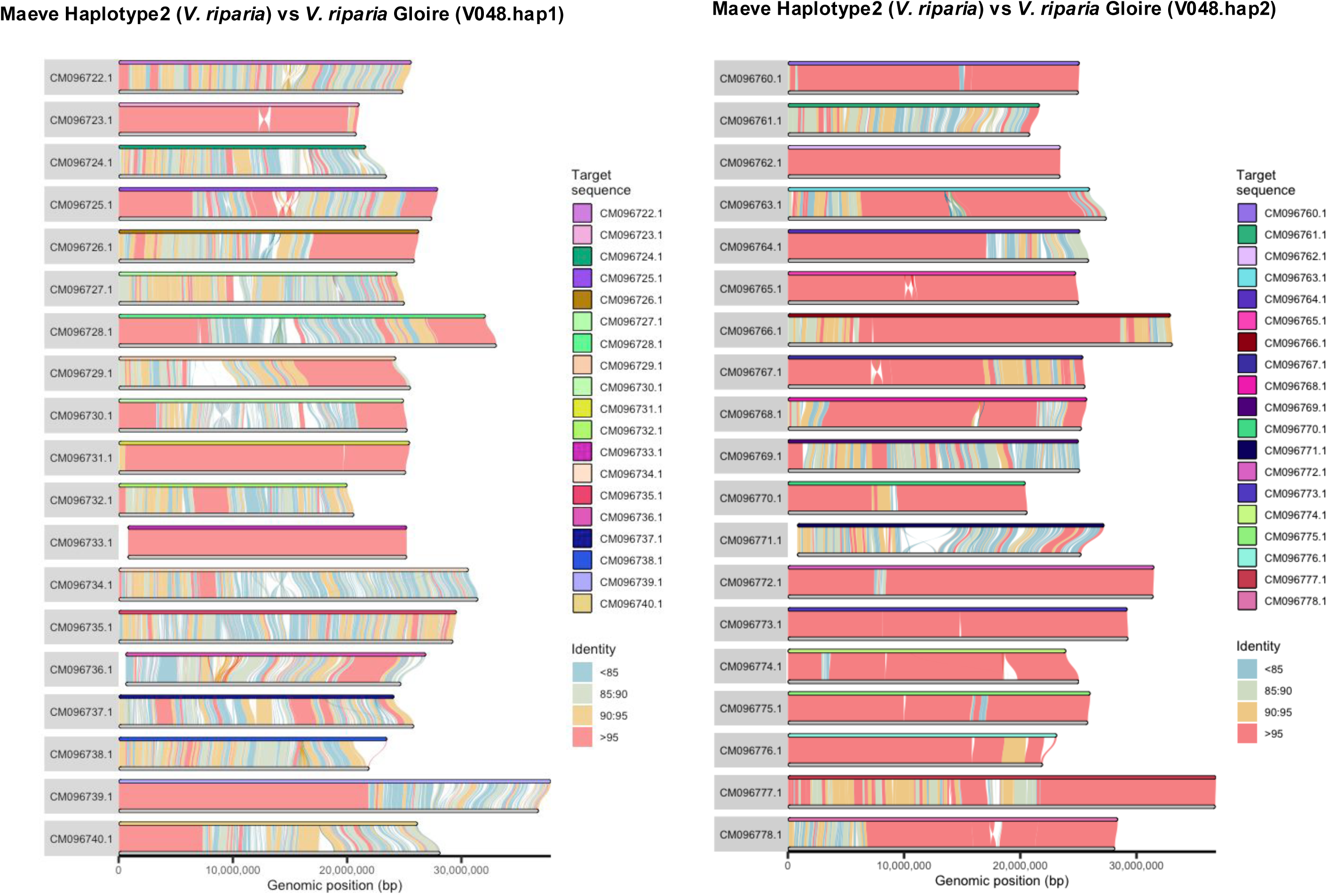
Synteny of Maeve haplotype 2 and the Vitis riparia Gloire genomes. Synteny blocks in chromosomes of *V. riparia* Gloire and haplotype 1 of the Maeve genome.

